# Exploring the variance in complex traits captured by DNA methylation assays

**DOI:** 10.1101/2020.10.09.333542

**Authors:** Thomas Battram, Tom R. Gaunt, Doug Speed, Nicholas J. Timpson, Gibran Hemani

## Abstract

Following years of epigenome-wide association studies (EWAS), traits analysed to date tend to yield few associations. Reinforcing this observation, we conducted EWAS on 400 traits and 16 yielded at least one association at the conventional significance threshold (P<1×10^−7^). To investigate why EWAS yield is low, we formally estimated the proportion of phenotypic variation captured by 421,693 blood derived DNA methylation markers (h^2^_EWAS_) across all 400 traits. The mean h^2^_EWAS_ was zero, with evidence for regular cigarette smoking exhibiting the largest association with all markers (h^2^_EWAS_=0.42) and the only one surpassing a false discovery rate < 0.1. Though underpowered to determine the h^2^_EWAS_ value for any one trait, h^2^_EWAS_ was predictive of the number of EWAS hits across the traits analysed (AUC=0.7). Modelling the contributions of the methylome on a per-site versus a per-region basis gave varied h^2^_EWAS_ estimates (r=0.47) but neither approach obtained substantially higher model fits across all traits. Our analysis indicates that most complex traits do not heavily associate with markers commonly measured in EWAS within blood. However, it is likely DNA methylation does capture variation in some traits and h^2^_EWAS_ may be a reasonable way to prioritise traits that are likely to yield associations.

## INTRODUCTION

Epigenome-wide association studies (EWAS) aim to assess the association between phenotypes of interest and DNA methylation across hundreds of thousands of CpG sites throughout the genome (1, 2). Many recent EWAS yielded few sites across the genome with strong evidence for association and the proportion of total trait variance associated with these sites is small (1). There is a need to have a global view of the contribution of DNA methylation to complex traits in order to interpret these results.

There are multiple possible reasons for there being few EWAS signals. Firstly, DNA methylation varies between cells and tissues, thus any changes related to a trait may occur in any number of tissues. Currently, because of ease of access and cost, the most common tissue used for EWAS is blood, which may not capture changes in DNA methylation related to the trait of interest (1, 2). Secondly, the commonly used technologies probe a small percentage of the total number of potentially methylated sites. Without knowing the full correlation structure across methylation sites, it is difficult to understand the coverage of current measures. Two more possibilities are that DNA methylation variation is actually not associated with the traits studied or that the associations are many but individually too small to detect with current sample sizes (Box 1).

Interpretation of the paucity of EWAS hits is difficult because there is no knowledge of the total contribution of methylation variation to the trait. However, analogous to the calculation of genetic heritability estimates, which have now been expanded to make inference across non-familial population-level data (SNP heritability), the total contribution of methylation markers to complex traits can potentially be estimated. This could give insight into the underlying patterns of association between DNA methylation markers and complex traits (See Box 2 for a simple explanation of SNP heritability (or h^2^_SNP_) and its application to DNA methylation (h^2^_EWAS_).

SNP heritability estimates are sensitive to assumptions of the underlying genetic architecture and there are different ways in which to model the contribution of each SNP to the overall genetic component. The original model of calculating h^2^_SNP_ introduced by Yang et al. assumes that each variant has an effect that is independent of the regional linkage disequilibrium (LD) structure as each variant is unweighted (the blanket model), and this effectively assumes regions of high LD contribute more to phenotypic variance (3). Speed et al. proposed a new model, which considered the LD between SNPs so that each region of high LD can effectively be counted as a singular effect (the grouping model)(4). Finding which models fit the data better helps ensure a more accurate estimation of the proportion of DNA methylation association with a trait, further, contrasting these models could also be biologically informative.

Gene regions are methylated in a coordinated fashion, which is associated with changes in gene expression (5, 6), with a tendency for promotor regions to be unmethylated and gene body regions to be methylated when gene expression is active (6). This, amongst other complex patterns of gene regulation, induces a correlation structure within EWAS data, and it is not clear whether a single site is driving an association and neighbouring sites are consequentially correlated, or if the cumulative contributions of all neighbouring sites associate with the regulatory process. In EWAS, a common strategy is to collapse DNA methylation sites into groups based on proximity and if they share the same direction of association and potentially magnitude of association, this is often called differentially methylated region (DMR) analysis (7). This, however, does not explain whether the sites within groups are acting independently and cumulatively or as a set of distinct influences. Figure 1 shows a representation of how the differences in models apply to DNA methylation data at a single small region in one specific example. Of course, there are far more scenarios possible and furthermore, the models aren’t restricted to a single small region in the genome. They apply to all sites, as do the DMR methods used in EWAS. Thus, by applying both methods to DNA methylation data across multiple phenotypes and comparing their utility we can gain insight into how DNA methylation operates across gene regions. Furthermore, it is important to find the model that best fits the data to help reduce bias in the estimates.

**Figure 1.**
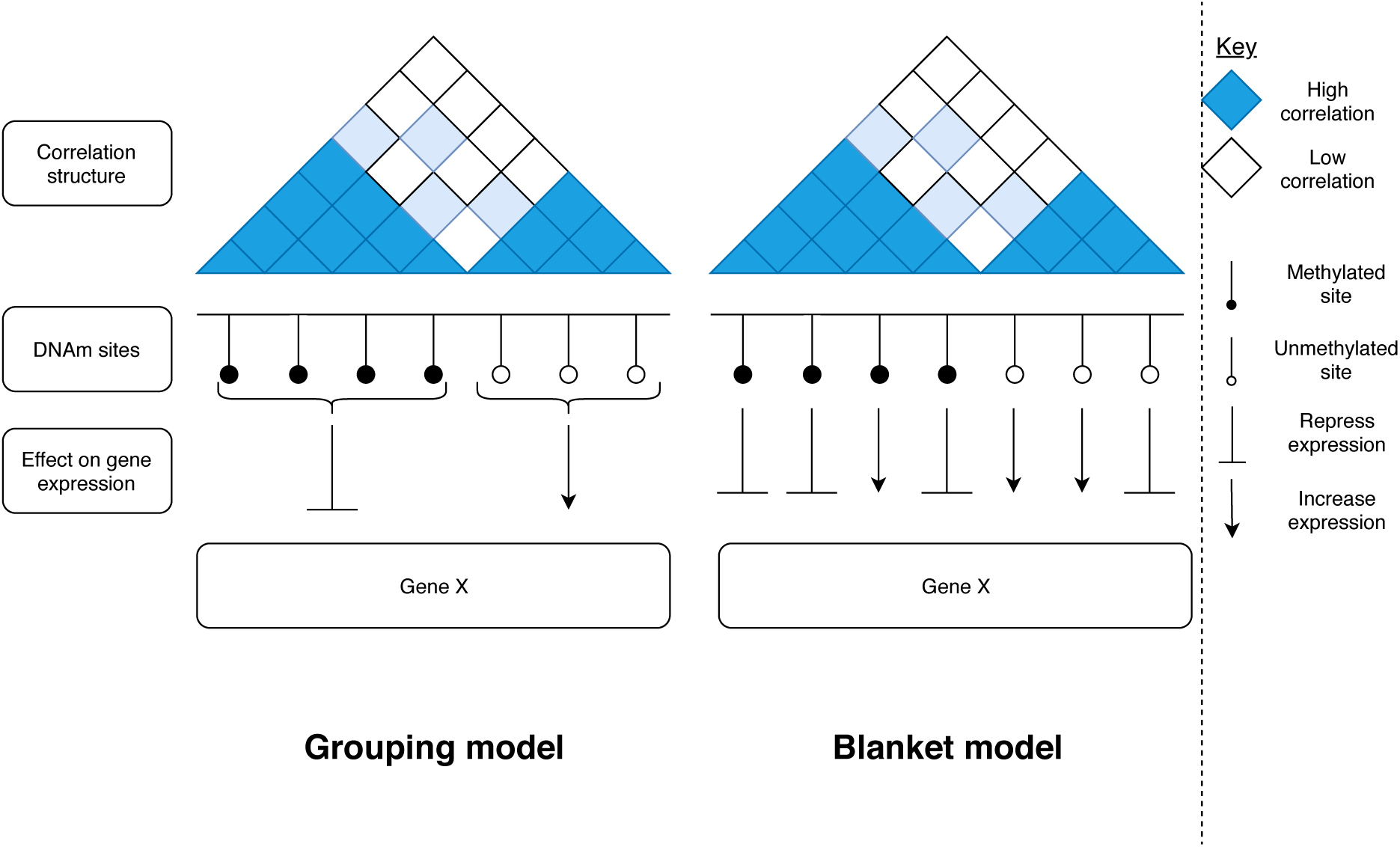
Comparison of the grouping and blanket models in the context of the relationship between DNA methylation and gene expression. Both regions are exactly the same, the only difference is how each model assumes the methylation sites should be treated. The grouping model down-weights the contribution of correlated CpGs, effectively grouping them, and the blanket model assumes each CpG independently associates with a trait. As seen here, the grouping of correlated CpG sites may not be the correct thing to do as some of the sites may be acting independently of their correlated partners.

This study aims to estimate h^2^_EWAS_ values across a plethora of traits and assesses whether this estimate may be useful in identifying traits for which EWAS will likely yield successful identification of associated DNA methylation sites. To do this we perform hundreds of EWAS studies and evaluate if h^2^_EWAS_ estimates are predictive of the number of sites identified by the EWAS at various P value thresholds. We also compare the performance of different models underlying h^2^_EWAS_ estimates to infer likely methylation architecture of complex traits.

### Box1: The argument for increasing sample size for EWAS

The need for larger sample sizes in GWAS has been empirically demonstrated across a broad range of traits. For height and body mass index (BMI), the number of associations dramatically increased from 12 to 3290 and from one to 941, respectively after increasing sample sizes by ~670,000 (27–29). This trend can be seen for many traits. Similar to early GWAS, many EWAS are discovering few sites strongly associated with complex traits. However, an example that suggests promise for increasing sample sizes for EWAS is seen with BMI, where an EWAS of 459 individuals identified just five sites, but increasing the sample size to over 5,000 led to identification of 278 sites (30, 31). While we can continue to improve sample sizes in EWAS, there is a need to determine the upper limit of the information we can obtain from EWAS of complex traits like BMI. Furthermore, the BMI EWAS example may be unrepresentative of other traits, so having a corollary test for estimating the contribution of common genetic variants to trait variance (h^2^_SNP_) for DNA methylation would help us understand if we’re capturing relevant information from the current arrays we are using in EWAS. Such information could inform future study designs in terms of growing sample sizes with the current assays available versus designing new assays.

### Box2: Applying SNP heritability estimator methods to DNA methylation

Methods used to estimate h^2^_SNP_ use restricted maximum likelihood (REML) tests to estimate the proportion of variance attributable to these genetic variants. Essentially this assesses whether individuals that are genetically similar are more likely to be phenotypically similar. If those individuals that have a high genetic overlap tend to correlate strongly phenotypically compared to those that do not have high genetic overlap, then the phenotype of interest will have a high h^2^_SNP_. Unlike genetic variants, DNA methylation is responsive to the environment (1) and determining causal directionality between DNA methylation markers associated with traits is not trivial (32–34). Therefore, estimating the proportion of trait variation captured by DNA methylation variation (which will henceforth be denoted as h^2^_EWAS_) using the same techniques will ascertain effects going in both directions as well as associations due to confounding. The combination of these mechanisms may increase power to detect trait-DNA methylation association, and could be the reason that so many DNA methylation markers are found in small EWAS compared to similarly sized GWAS (31).

## MATERIAL AND METHODS

### Study samples

All data for the study came from the Avon Longitudinal Study of Parents and Children (ALSPAC) cohort. Pregnant women resident in Avon, UK with expected dates of delivery 1st April 1991 to 31st December 1992 were invited to take part in the study. The initial number of pregnancies enrolled is 14,541 (for these at least one questionnaire has been returned or a “Children in Focus” clinic had been attended by 19/07/1999). Of these initial pregnancies, there was a total of 14,676 foetuses, resulting in 14,062 live births and 13,988 children who were alive at 1 year of age. Full details of the cohort has been published previously (8, 9). This study uses phenotypic and DNA methylation data from the mothers (N = 940).

Continuous and binary phenotypes measured in mothers were extracted from the cohort. A summary of the phenotypes is present in the **Supplementary Material**. Please note that the study website contains details of all the data that is available through a fully searchable data dictionary and variable search tool http://www.bristol.ac.uk/alspac/researchers/our-data/

Phenotype data were extracted using the ‘alspac’ R package (github.com/explodecomputer/alspac) and went through various quality control steps, which are detailed in the **Supplementary Material** and summarized in **Supplementary Figure 1**.

All continuous traits were rank-normalised for further analyses. A Shapiro-Wilk test of normality was performed on these rank-normalised traits and for those with some evidence of non-normality (P < 0.05), we re-examined the distribution of those traits by eye to ensure it was approximately normal. It was found that any non-normality of phenotype distributions corresponded to an inflation of zero values. These traits were removed and overall there were 2408 traits left for analyses. These traits do not necessarily represent independent phenotypes and as such we wanted to prevent correlated traits skewing results. The absolute Pearson’s correlation coefficient between each trait was subtracted from one (1 –[r]). Then traits were greedily selected where 1 –[r] < 0.4 with any other trait. This left 400 traits, which consisted of ~30% clinically measured variables (including roughly 50 metabolites and some anthropometric traits), ~25% health related questions (for example “have you ever had asthma?”), ~40% behavioural and social traits (for example educational attainment variables, use of pesticide, and having pets), and ~5% of traits were related to the partner or child of the participant (for example the employment status of the partner). Phenotypes are presented in **Supplementary table 1**. Plots showing the correlation between all the phenotypes as well as with just the selected traits can be seen in **Supplementary Figure 2-3**.

Ethical approval for ALSPAC was obtained from the ALSPAC Ethics and Law Committee and from the UK National Health Service Local Research Ethics Committees. Written informed consent was obtained from both the parent/guardian and, after the age of 16, children provided written assent. Consent for biological samples has been collected in accordance with the Human Tissue Act (2004). Informed consent for the use of data collected via questionnaires and clinics was obtained from participants following the recommendations of the ALSPAC Ethics and Law Committee at the time.

### DNA methylation data

DNA methylation was measured using the Illumina Infinium HumanMethylation450 (HM450) BeadChip. Before use, the data went through quality control and were normalised separately to the phenotype data. Full details can be found in the **Supplementary Material**.

DNA methylation data generated from blood collected at a single clinic visit was used for each of the participants.

Probes were excluded if they were present on either of the sex chromosomes, a SNP/control probe, had a detection p value < 0.05 across over 10% of samples or were identified as problematic by Zhou et al. (10). This left 421,693 CpG sites for analyses.

Before analysis a linear regression model was fitted with beta values for methylation (which ranges from 0 (no cytosines methylated) to 1 (all cytosines methylated)) as the outcome against batch variables (plate ID in ALSPAC) modelled as a random effect to help remove the effects of batch in the subsequent analyses.

Cell proportions (CD8+ and CD4+ T cells, B cells, monocytes, natural killer cells, and granulocytes) were estimated using an algorithm proposed by Houseman et al. (11).

### REML analysis

Using LDAK (12) the relationship between the methylomes (as measured by the HM450 BeadChip) of 940 individuals was estimated by producing a DNA methylation relationship matrix (MRM). This matrix was used as input for the REML analysis to estimate the proportion of a trait’s variation that was correlated with DNA methylation (h^2^_EWAS_). Age, the top 10 ancestry principal components, and derived cell proportions were added as covariates to the model.

When producing the MRM, probes were scaled by their observed variance and the weighting of each probe was based on the variance of DNA methylation at that site using the formula below:

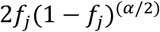

where *f*_*j*_(1 − *f*_*j*_) is the variance of methylation at CpG *j*.

The higher the alpha value the more weight is given to probes with greater variance; an alpha value of −1 gives equal weight to probes with low and high variance. The alpha value of −0.25 was chosen because previous analysis by Speed et al. (12) suggested that this value was optimal for measuring h^2^_SNP_. Furthermore, it was hypothesised that probes with a greater variance would contribute more to trait variance. As the method was applied to DNA methylation data in this study, sensitivity analyses were conducted. MRMs were created specifying the alpha value at increasing increments of 0.25 from −2 to 0. The variation of h^2^_EWAS_ and how well the model fit the data was assessed for the varying alpha values.

The mean of the MRM diagonal should be 1 and the variance close to 0, as the diagonal values essentially represent the correlation between an individual’s methylome with itself. Although values are expected to vary slightly from 1. For the MRMs it was identified that some diagonal elements were very high (> 2), which caused the diagonal to have a high variance (0.13). To assess whether these values could skew results, we conducted sensitivity analyses removing individuals, with varying diagonal value cut-offs.

Like h^2^_SNP_ estimates, h^2^_EWAS_ estimates should range from zero to one. If a trait has a true h^2^_EWAS_ value of zero, there is no association between the methylome and that trait, and if h^2^_EWAS_ equals one then DNA methylation has the capacity to completely predict that trait.

However, estimation of h^2^_EWAS_ can be fairly imprecise and without constraining the software it’s possible to get estimates of h^2^_EWAS_ that are outside 0-1 due to large standard errors. These point estimates have to be erroneous by definition.

Even though the grouping model effectively groups sites together, it is actually likely to increase the number of parameters because without the weightings imposed by this model, the blanket model essentially ignores sites that are not neighbouring others. Therefore, larger standard errors are expected with the grouping model. The grouping model applies a sliding window approach, with windows of 100kb, to capture the correlation between neighbouring sites and weight sites according to the correlation structure of the region. When applying the grouping model, the number of sites that were weighted were 45,863 (out of 421,693) and the number of sites neighbouring any single CpG site ranged from 29 to 28,217.

### Generating genetic principal components

Ancestry principal components were generated within ALSPAC mothers using PLINK (v1.9). Before analysis, genetic data went through quality control and were imputed, full details can be found in the **Supplementary Material**. After quality control and imputation, independent SNPs (r^2^ < 0.01) were used to calculate the top 10 ancestry principal components.

### Epigenome-wide association studies

EWAS were conducted for 400 selected traits (see **Study samples** section for selection process) within the ALSPAC cohort. For all traits, linear regression models were fitted with beta values of DNA methylation as the outcome and the phenotype as the exposure. Covariates included age, the top 10 ancestry principal components and cell proportions.

### Association between h^2^_EWAS_ and epigenome-wide association studies results

Differentially methylated positions (DMPs) were extracted from the EWAS at P value thresholds ranging from 10^−3^ to 10^−7^. It was assessed whether h^2^_EWAS_ could predict that the number of identified DMPs in an EWAS was greater than number of DMPs expected to be identified at a given P threshold defined as the number of sites tested multiplied by the threshold. The traits were also “pruned” in the same way as described above, to prevent including overly correlated traits and biasing results. The sensitivity and specificity of this prediction was calculated and a receiver operating characteristic (ROC) curve was plotted. At p-value thresholds of 10^−6.5^ and 10^−7^ there were too few EWAS hits, so these were removed from the analysis.

The association between the number of DMPs identified at P < 1×10^−5^ and h^2^_EWAS_ values was assessed using a negative binomial hurdle model with the number of DMPs identified fitted as the outcome and h^2^_EWAS_ as the exposure. The negative binomial hurdle Poisson regression model results are twofold. The first of which assesses whether there is an association between the binary trait of whether a DMP was identified by EWAS and h^2^_EWAS_. The second is a zero-truncated model, i.e. the zero values are removed from the model and the association between number of DMPs and h^2^_EWAS_ is assessed.

The same method was applied to estimate the association between the number of SNPs identified in GWAS at P < 5×10^−8^ and h^2^_SNP_. SNPs associated with 485 traits in UK Biobank (see **Supplementary Material** for sample information and phenotype selection) were extracted using the IEU Open GWAS Database (13). The h^2^_SNP_ estimates were extracted from http://www.nealelab.is/uk-biobank/.

All analyses were conducted in R (version 3.3.3) or using the command line software LDAK (12), GCTA (14), and PLINK (15). For the EWAS analyses, the meffil R package was used (16). A one-sided P value was used to assess if the h^2^_EWAS_ for a trait was > 0, and two-sided P values were used for everything else.

## RESULTS

A flowchart showing our study design and giving a summary of the results is shown in Figure 2.

**Figure 2.**
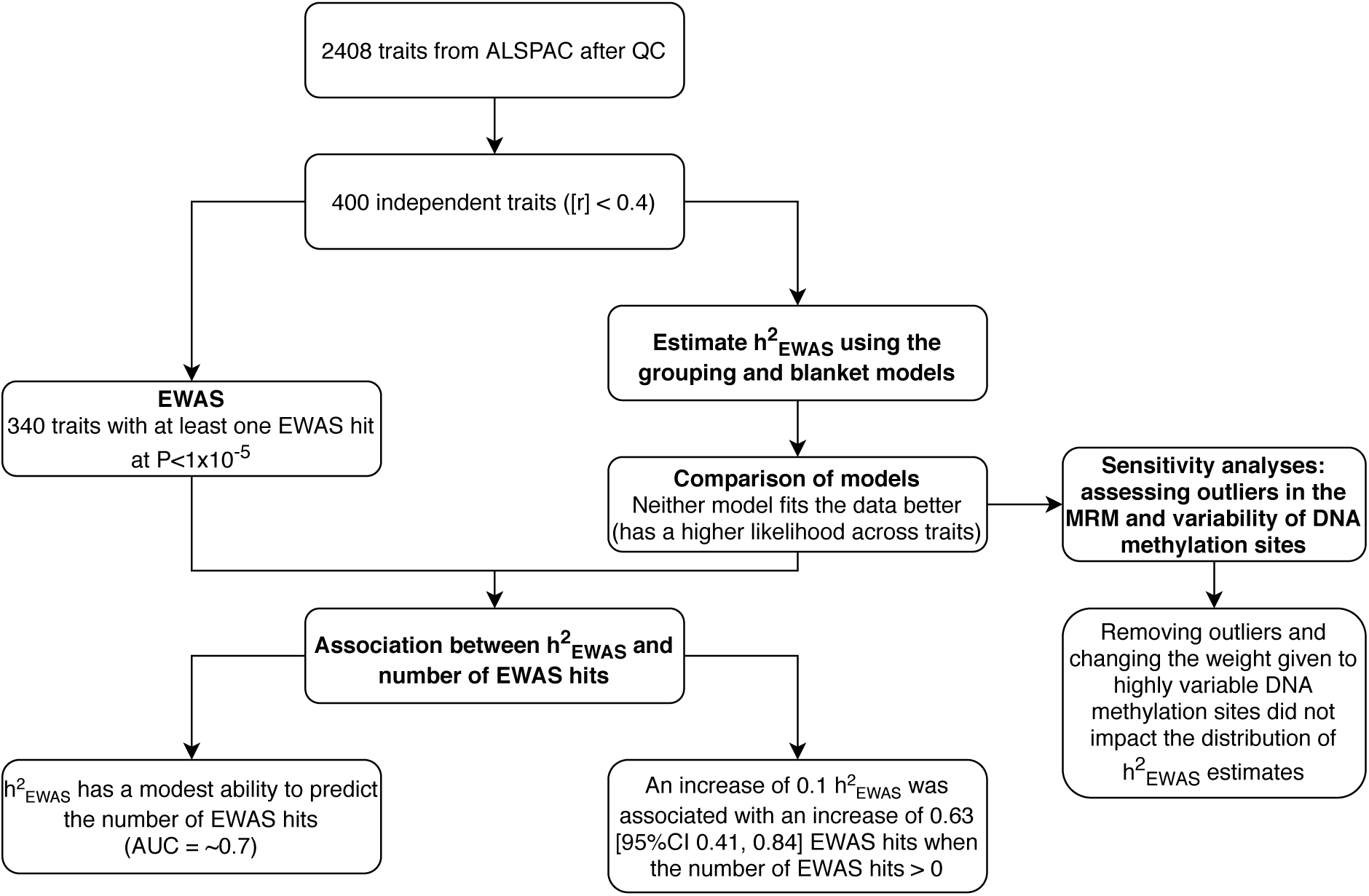
Study design with a summary of the results. ALSPAC = Avon Longitudinal Study of Parents and Children, QC = quality control, EWAS = epigenome-wide association study, MRM = methylation relationship matrix, AUC = area under curve.

### Estimating the proportion of phenotypic variance associated with DNA methylation

We used two models to estimate the total contribution of all DNA methylation sites to the variation (h^2^_EWAS_) for each of 400 traits within 940 individuals. The mean for both models was zero with ranges of −0.4 to 0.4 and −0.5 to 0.4 for the blanket and grouping models respectively Figure 3. The estimates were imprecise, the mean standard error was 0.03 and 0.05 for the blanket and grouping models respectively. The trait with the greatest evidence for h^2^_EWAS_ estimates being above zero was having smoked cigarettes regularly (FDR-corrected P = 0.06 and 0.10 for the blanket and grouping models respectively). The correlation between the h^2^_EWAS_ estimates of the two models was 0.47 and there was evidence that on average the estimates of the grouping model were higher (Paired t-test P = 1.8×10^−5^, Figure 3), but the mean difference between estimates was only 0.018.

**Figure 3.**
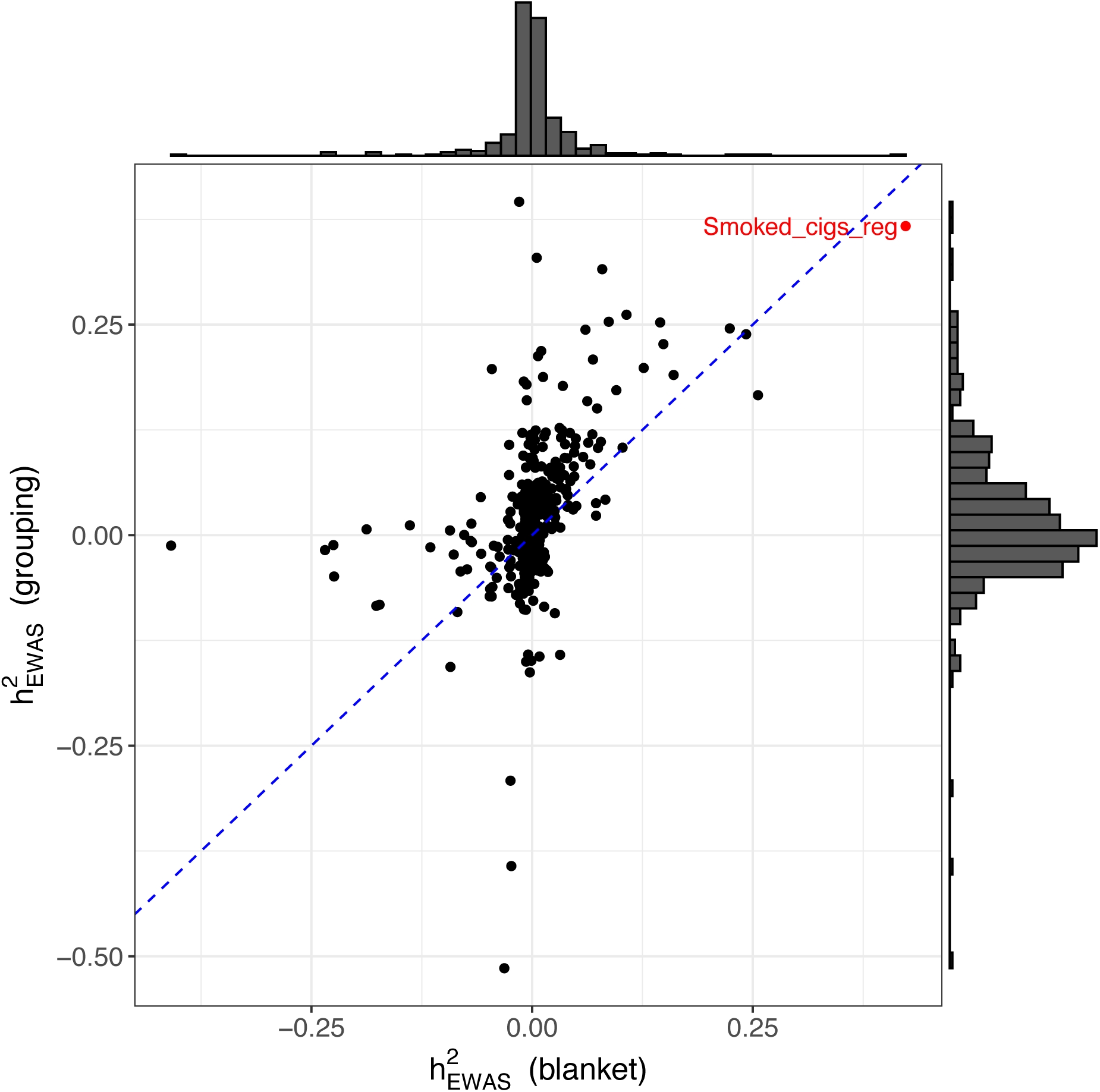
A comparison of h^2^_EWAS_ estimates given by applying REML using the blanket and grouping models across 400 traits. The blue dashed line is at x=y. Values with h^2^_EWAS_ lower than 0 are due to imprecision in h^2^_EWAS_ estimates as the true estimate cannot be negative. Smoked_cigs_reg = smoked cigarettes regularly. The h^2^_EWAS_ of this phenotype has the greatest evidence for being above 0 for both the blanket and grouping model (Uncorrected P = 1.44×10^−4^ and P = 2.61×10^−4^, respectively).

There was little evidence that either of the models fit the data better (had higher likelihoods) across the 400 traits tested (difference in median likelihoods = 0.19, Wilcoxon’s paired ranked sum test P = 0.73). Further, the majority of h^2^_EWAS_ estimate differences between the traits were small.

### Sensitivity analyses when estimating the proportion of phenotypic variance associated with DNA methylation

After examination of the MRMs required to produce the h^2^_EWAS_ estimates, we found that for both the blanket and grouping model we observed some unexpected values: 96 diagonal elements had values over 1.5 when using the blanket model, with the maximum value being 3.562. When assessing the impact of these potential outliers in the MRM to results we found that the median and range of h^2^_EWAS_ estimates varied little (**Supplementary Figure 4**). The likelihood of the models tended to be greater as more outliers were removed (lower threshold for classing a diagonal element as an outlier), but it still didn’t vary much (**Supplementary Figure 5**).

The weight of predictors used to produce the MRMs was also examined. As more weight was given to sites where methylation variation was greater (increasing alpha value) the h^2^_EWAS_ estimates were slightly higher (**Supplementary figure 6**). However, the likelihood tended to remain the same, the median likelihood had a range of 2 across the alpha values (**Supplementary figure 7**).

Results of sensitivity analyses are summarised in **Supplementary table 2 and 3**.

### EWAS analyses

In order to assess the association between h^2^_EWAS_ and EWAS results, we performed EWAS of 400 traits. No associations were found at the strict P value cutoff of P < 2.5×10^−10^ (conventional EWAS P-value threshold, 1×10^−7^, divided by the number of traits, 400). A total of 29 associations between traits and CpGs were identified at the conventional EWAS P value cutoff – P < 1×10^−7^. Of the traits tested, 16 had at least one EWAS hit, with the maximum number of CpGs associated with a trait being 13 (smoked cigarettes regularly). As there were so few traits with any identified hits, we took forward results from the lenient P value threshold of P < 1×10^−5^, at which 340 traits had at least one EWAS hit. **Supplementary table 4** shows each trait and the number of DMPs identified at varying p-value thresholds.

The distribution of the number of DMPs identified was heavily right skewed with an inflation at 0 and 1 (**Supplementary figure 8**), therefore, to test the association between h^2^_EWAS_ and number of DMPs we opted to test goodness of fit for variations of Poisson models. Of the 6 models tested, the negative binomial hurdle Poisson regression model fit the data best, full results can be found in **Supplementary table 5**. We found there was some evidence for an association between number of DMPs identified and h^2^_EWAS_ (Figure 4). There was some evidence of association between the presence of DMPs and h^2^_EWAS_ (beta = 6.2, [95%CI 2.5, 10]) as well as some evidence of an association between number of DMPs (when the number is above 0) and h^2^_EWAS_ (mean increase of 0.63, [95%CI 0.41, 0.84] DMPs when h^2^_EWAS_ increases by 0.1). Applying the same method to GWAS data we found evidence that the presence of identified SNPs associated with h^2^_SNP_ (beta = 21.9 [95%CI 19.6, 24.1]) and the association between number of SNPs identified (when the number is above 0) and h^2^_SNP_ (mean increase of 1.5, [95%CI 0.93, 2.5] SNPs when h^2^_SNP_ increases by 0.1).

The ability of h^2^_EWAS_ estimated by both models to predict whether the number of DMPs identified was greater than expected was assessed at varying P value thresholds. ROC curves were produced and the area under the curve (AUC) ranged from 0.65 and 0.67 at P < 1e-6 to 0.79 and 0.71 at P < 1e-3 for the blanket and grouping models respectively and the predictive ability remained fairly stable as the threshold increased (Figure 5).

**Figure 4.**
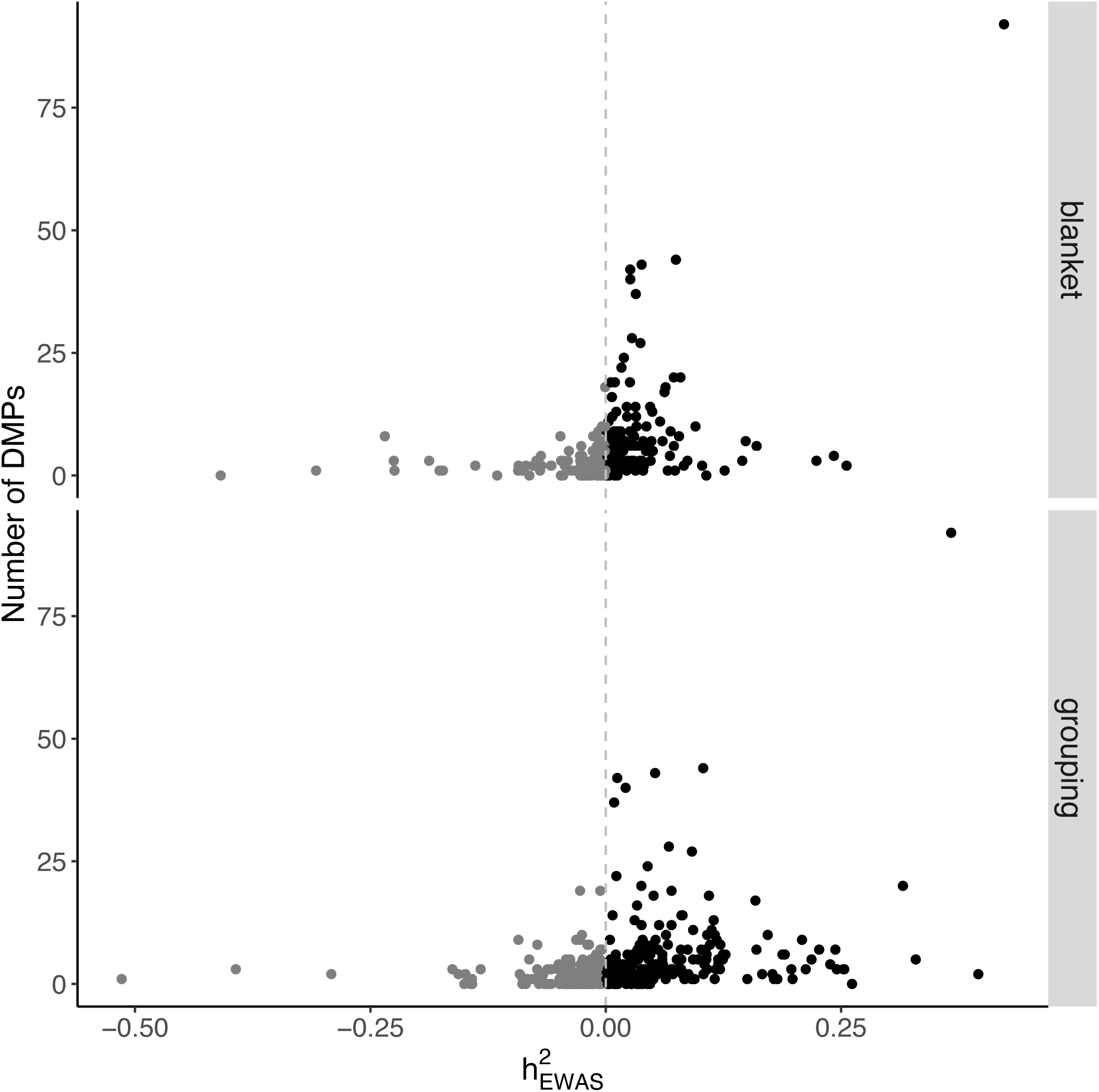
Association between h^2^_EWAS_ and number of DMPs identified in EWAS. The correlation between DNA methylation and the variance of traits (h^2^_EWAS_) was calculated using REML analysis using the blanket and grouping models. EWAS were conducted on all the same traits and the distribution of the number of DMPs identified at P < 1×10^−5^ and h^2^_EWAS_ are plotted above. Any traits where the h^2^_EWAS_ estimate is below 0 are coloured grey. The true h^2^_EWAS_ value of a trait cannot be negative, but sample sizes in this analysis are small so the estimates are imprecise.

**Figure 5.**
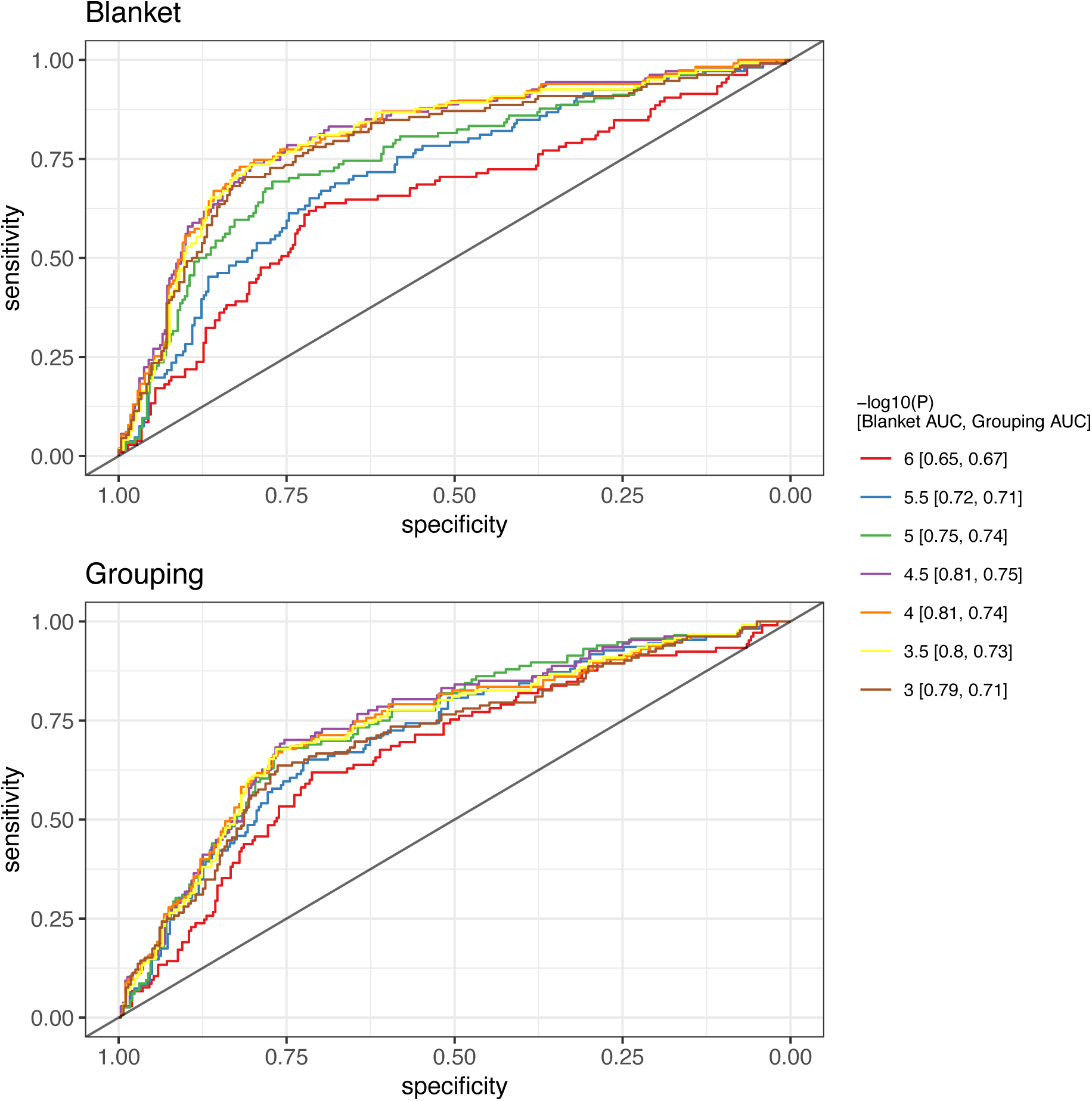
The ability of h^2^_EWAS_ values to predict whether the number of differentially methylated positions identified in an EWAS is higher than expected by chance. ROC curves for h^2^_EWAS_ values predicting number of DMPs at differing P value thresholds. AUC = area under the curve.

## DISCUSSION

The global contribution of DNA methylation to complex trait variance can inform researchers of how to design future studies that seek to discover new DNA methylation sites associated with their trait of interest. In this manuscript we apply methods designed to estimate the predictive capacity of variants across a SNP-chip (h^2^_SNP_), to DNA methylation data measured in blood with the HM450 BeadChip across 400 independent traits, giving a distribution of the contribution of all sites typically measured in EWAS to complex trait variance. Although sample size was too small to reliably estimate h^2^_EWAS_ for any one trait, the distribution of estimates suggest little complex trait variation is captured by DNA methylation at the sites measured and h^2^_EWAS_ may be a good measure to identify traits for which EWAS will yield associations.

### Estimation of h^2^_EWAS_

The true h^2^_EWAS_ of a trait gives the total predictive capacity of DNA methylation for that trait, which is equivalent to the proportion of that trait’s total variance that is associated with changes in DNA methylation. Knowing this information can help design future EWAS studies. A low value of h^2^_EWAS_ doesn’t necessarily mean there is little correlation between DNA methylation and a trait, it could transpire that unmeasured sites contribute more to the association. It is important to remember that roughly 1.5% of CpG sites are targeted by the HM450 BeadChip and DNA can be methylated elsewhere (not at cytosine bases). Therefore, whole genome bisulphite sequencing, or a similar technique, may show that the variance of complex traits captured by DNA methylation is far higher. Furthermore, even if h^2^_EWAS_ is low and the sites discovered already do not explain all of the h^2^_EWAS_ estimate, there may still be value in increasing sample size to identify more DMPs as well as increase the precision of h^2^_EWAS_ estimates. DMPs discovered may not be highly correlated with a trait, but the potential biological information gained may still be valuable. For example, if a change in the levels of protein X has a large effect on a trait and change in DNA methylation has a small effect on the levels of protein X, then the effect of that DNA methylation change on the trait will be small, but identifying that DMP could lead to discovering the importance of the protein. Another thing to consider is that DNA methylation is tissue and cell specific. This means, that h^2^_EWAS_ may vary a lot depending on what tissue the methylation is measured in.

The true underlying genetic architecture of complex traits is still unknown, and therefore it is difficult to know the appropriate model to choose when estimating the contribution of all measured SNPs to phenotypic variance amongst unrelated individuals and arguments for each model depending on this underlying genetic architecture are still being put forward (12, 17–19). Thus, the attempts made in this study to re-purpose genomic REML are likely to suffer the same flaws that are trying to be overcome in genetic data. With this in mind, in addition to the imprecise estimates of h^2^_EWAS_ presented here (due to the small sample sizes of available data), we believe that individual trait h^2^_EWAS_ values should be treated with caution. This doesn’t exclude the possibility that estimating h^2^_EWAS_ may be useful and other methods are already being developed to measure the association between DNA methylation at all sites and complex traits (20).

### Future EWAS

Heritability estimates from family-based studies gave an *a priori* justification for the pursuit of gene mapping endeavours that eventually gave rise to GWAS, as they demonstrated variation in complex traits had a substantial genetic component. However, the evidence DNA methylation contributed to trait variation was not ascertained before EWAS were first conducted. To justify collecting more samples and continuing with EWAS research in the current vein, methods such as the one presented in this study should be used to show DNA methylation does substantially contribute to trait variance. It has become clear from the GWAS era of genetics, that for complex traits, such as coronary artery disease, many common genetic variants with small effects make up the genetic component of the trait (21, 22). This suggests a large number of molecular pathways contribute to these traits. DNA methylation at CpGs is heritable (23, 24), thus it would be expected that the DNA methylation architecture of a trait will somewhat reflect the genetic architecture of the trait, although this has not been empirically tested.

Despite uncertainty of h^2^_EWAS_ estimates for individual traits, we show h^2^_EWAS_ has a modest ability to predict whether the number of EWAS hits will be greater than expected by chance at a given P value threshold. This predictive ability remained stable as the P value threshold for detection increased from P < 1×10^−6^ to P < 1×10^−3^. These results suggest that increasing sample sizes for traits which truly associate with DNA methylation should result in the discovery of more DMPs. Furthermore, these results support a model for which small changes in methylation at many CpGs across the genome are related to complex traits.

### Contributions of individual CpG sites

The original model for measuring h^2^_SNP_ assumed all genetic variants contributed the same effect on a trait (3), Speed et al. offered an alternative model assuming a different underlying genetic architecture, whereby genetic variants in regions of high LD contributed less to the variance of a trait than more independent variants. Both groups have shown that the performance of the models depend on the alignment of the trait’s architecture with the models’ underlying assumptions. Previous literature has suggested that it is the methylation across groups of CpGs that may affect how other molecules interact with DNA and influence cellular functions such as gene expression (6). Furthermore, CpGs are not randomly distributed throughout the genome – many exist in close proximity within “islands” or other regions, suggesting that grouping of the CpGs may have functionality. However, the most common method used in EWAS is to treat CpG sites as independent. Here, the models proposed by Speed et al. (the grouping model) and Yang et al. (the blanket model), when estimating h^2^_EWAS_ were tested across 400 traits. The model fit the data better (had a higher likelihood) 207 times for the blanket model and 193 times for the grouping model. Thus for almost half the traits treating DNA methylation sites as independent seems to be preferable and even though there is correlation between CpG sites, which allows them to be grouped, it might be that in some groups of correlated sites, individual sites within the group contribute separately to trait variance. It’s important to note that the grouping method takes into account correlation between CpGs within 100Kb of each other. Differential methylation at CpG sites may be correlated for a variety of biological reasons, for example, CpGs lying within a transcription factor binding site will be regulated together, but also, they will be correlated with CpGs that lie in other binding sites for that same transcription factor and these may be many megabases away. This is relevant to the relationship between DNA methylation and complex traits because transcription factor regulation might be the link between complex traits and DNA methylation. Even though grouping CpG sites might yet be the best way to model the relationship between DNA methylation and complex traits, the optimum way to group sites is unknown and will likely change depending on the trait of interest.

### Limitations

The main limitation of the study is the small sample size (N = 940) to estimate the h^2^_EWAS_. This meant the precision of our h^2^_EWAS_ estimates were very low and so our power to assess their ability to predict number of DMPs and find individual trait h^2^_EWAS_ values was low. To circumvent this problem, we assess trends across multiple traits and do not make strong conclusions for any one trait. As mentioned previously the HM450 BeadChip captures a small percentage of the total DNA methylome and h^2^_EWAS_ estimates will likely vary upon assaying more DNA methylation sites. Furthermore, when measuring more sites, it might be that one of the models fits the data better. Nevertheless, the results of this study can still give evidence towards the hypothesis that differential methylation at many sites across the genome each contribute minimally to the overall association between the methylome and a complex trait.

Unlike germline genetic variants, there is intra-individual (between tissue and over time) DNA methylation variation (1, 2). Thus, it is to be expected that the variation of h^2^_EWAS_ estimates across traits is partly a product of the tissue and timepoint of choice. However, within the tissue biologically pertinent to the complex trait of interest, the number of pathways that associate with variation in that trait is likely to remain high, for example there are many processes affecting, and affected by, cancer development (25). Thus, it would still be expected that differential methylation at many CpG sites each associate with a trait, but the effect sizes are small. The same can be said when estimating h^2^_EWAS_ at various timepoints.

Estimates of h^2^_EWAS_ will be a product of their environment and genetic makeup of the participants it’s measured in. Therefore, the results here may vary by population and by sex. However, participants used in this study are considered to be representative of the larger ALSPAC cohort (26), which is itself considered to be representative of a large majority of women from the UK and potentially other high-income countries (8). This suggests the results will be generalisable to a large group of samples for which EWAS are conducted, but replication in these samples as well as in different populations would provide greater confidence in the generalisability of the results.

A wide range of complex traits was used in the analysis, but there are some notable absences. Rarer diseases and diseases that predominantly impact the elderly are not present in this study. The results presented here cannot be generalised to those traits.

The factors important for the correlation structure of DNA methylation data are less known than those for linkage disequilibrium structure of genetic variants. Therefore, when applying models, such as the grouping model here, that aim to account for correlation of neighbouring DNA methylation sites, we may be missing some of the important structure captured for example by trans-correlations (over 1Mb). A model that estimates h^2^_EWAS_ by incorporating all of the underlying correlation of DNA methylation data may therefore outperform both models tested here.

### Conclusion

Overall, the number of traits with good evidence for h^2^_EWAS_ > 0 was low (only smoking behaviour met the threshold of FDR < 0.1) and mean h^2^_EWAS_ value across both models was roughly 0, suggesting that for many traits DNA methylation variation as measured on the HM450 BeadChip in blood is of little relevance. However, these estimates varied greatly and therefore DNA methylation measured in this way will likely have relevance for some traits, for example smoking cigarettes regularly. Further, these estimates were correlated with the number of DMPs identified, suggesting that for traits whose variance associates with DNA methylation then increasing sample size will yield an increase in the number of CpGs identified in EWAS. We also provide evidence that there is value in assessing individual CpG-trait associations as opposed to groups of correlated CpG sites within 100Kb. However, this does not preclude the possibility that a more complex model of CpG site correlation may provide a better fit.

## AVAILABILITY

Code used to perform analyses can be found here: https://github.com/thomasbattram/ereml

Data of the 400 EWAS performed are available at the University of Bristol data repository, data.bris, at https://doi.org/10.5523/bris.2bcpmkslk93a52gp8mb5tzh9j4

## SUPPLEMENTARY DATA

All supplementary figures and tables except supplementary table 1 and supplementary table 4 are part of the **Supplementary material**.

Supplementary table 1 and supplementary table 4 can be found in their own spreadsheet.

## ACKNOWLEDGEMENT

We are extremely grateful to all the families who took part in the Avon Longitudinal Study of Parents and Children the midwives for their help in recruiting them, and the whole ALSPAC team, which includes interviewers, computer and laboratory technicians, clerical workers, research scientists, volunteers, managers, receptionists and nurses.

