## Supplementary Material for "Exploring the variance in complex traits captured by DNA methylation assays"

### SUPPLEMENTARY METHODS

#### Phenotype data

The data that were extracted from ALSPAC had to go through extensive quality control. Originally over 15,000 traits that were related to the mothers were extracted from the database. After removing categorical variables, variables with >50% missing data and more, the final number of traits was 2408. A summary of this process is in **Supplementary Figure 1**. Traits were also removed if they were thought to have no relevance to the mother for example, the fieldworker that examined the mother several months or years after blood draw.

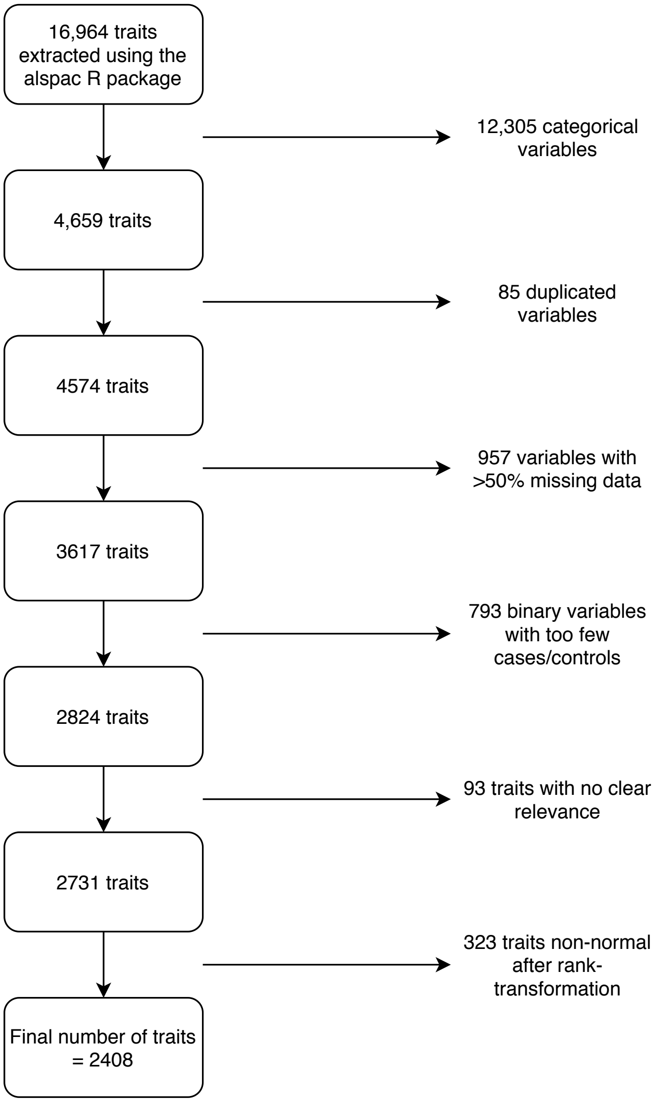


**Supplementary figure 1** A summary of the data cleaning steps.

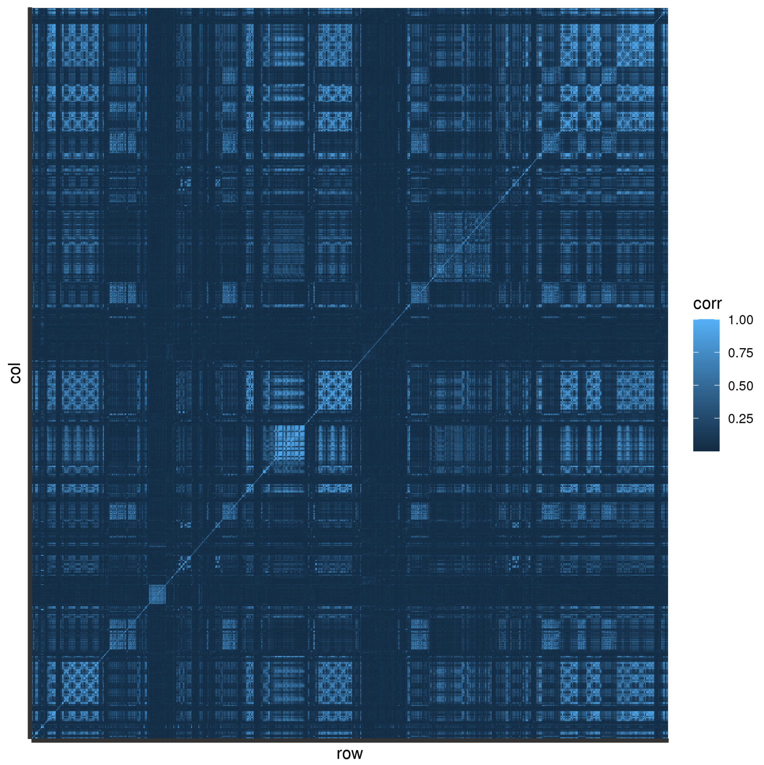


**Supplementary figure 2** Correlation between all 2408 phenotypes

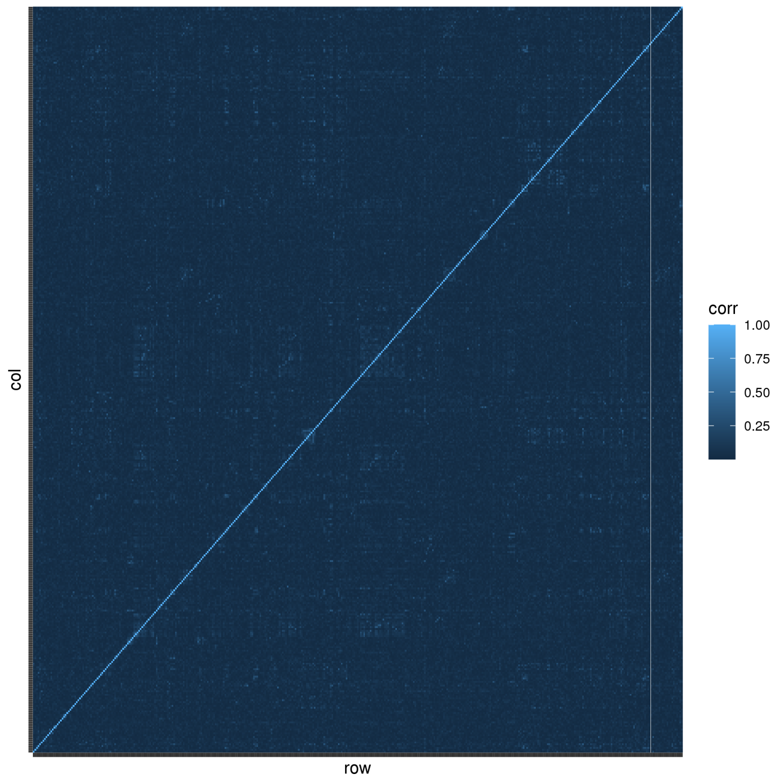


**Supplementary figure 3** Correlation between the 400 “independent” traits

### DNA methylation data

Following DNA extraction samples were bisulfite converted using the Zymo EZ DNA Methylation kit (Zymo, Irvine, CA). Genome-wide methylation was measured using the Illumina Infinium HumanMethylation450 (HM450) BeadChip. The arrays were scanned using an Illumina iScan, with initial quality review using GenomeStudio. During the data generation process, a wide range of batch variables were recorded in a purpose-built laboratory information management system (LIMS). The LIMS also reported quality control (QC) metrics from the standard control probes on the HM450 BeadChip for each sample. Methylation data were normalised in R with the wateRmelon package (1) using the Touleimat and Tost (2) algorithm to reduce the non-biological differences between probes.

### Genotyping

Mothers were genotyped using the Illumina human660W-quad genome-wide SNP genotyping platform (Illumina Inc., San Diego, CA, USA) at the Centre National de Génotypage (CNG; Paris, France). SNPs were removed if they displayed more than 5% missingness or a Hardy-Weinberg equilibrium P value of less than 1.0e-06. Additionally, SNPs with a minor allele frequency of less than 1% were removed. Samples were excluded if they displayed more than 5% missingness, had indeterminate X chromosome heterozygosity or extreme autosomal heterozygosity. Samples showing evidence of population stratification were identified by multidimensional scaling of genome-wide identity by state pairwise distances using the four HapMap populations as a reference, and then excluded. Cryptic relatedness was assessed using a IBD estimate of more than 0.125 which is expected to correspond to roughly 12.5% alleles shared IBD or a relatedness at the first cousin level. Related subjects that passed all other quality control thresholds were retained during subsequent phasing and imputation.

Imputation of mother's genotype data in ALSPAC was done with ALSPAC children's data. So, genotypes in common between the sample of mothers and sample of children were combined. SNPs with genotype missingness above 1% due to poor quality were removed along with subjects due to potential ID mismatches. We estimated haplotypes using ShapeIT (v2.r644) which utilises relatedness during phasing. We obtained a phased version of the 1000 genomes reference panel (Phase 1, Version 3) from the Impute2 reference data repository (phased using ShapeIt v2.r644, haplotype release date Dec 2013). Imputation of the target data was performed using Impute V2.2.2 against the reference panel (all polymorphic SNPs excluding singletons), using all 2186 reference haplotypes (including non-Europeans).

### UK Biobank

Over 500,000 people aged 37-73 years were recruited from the United Kingdom between 2006 and 2010 for the UK Biobank cohort. The study, participants, and quality control of phenotype and genotype data have been described previously (3–5).

GWAS conducted by the Neale Lab (http://www.nealelab.is/uk-biobank/) with sample size greater than 5000 were extracted for analysis. Any binary variables had to have more than 100 individuals per category and duplicated traits were removed from the analysis. Independent SNPs ($r^2 < 0.001$) were extracted using the 'extract_instruments' function from the TwoSampleMR package (6).

**SUPPLEMENTARY RESULTS**

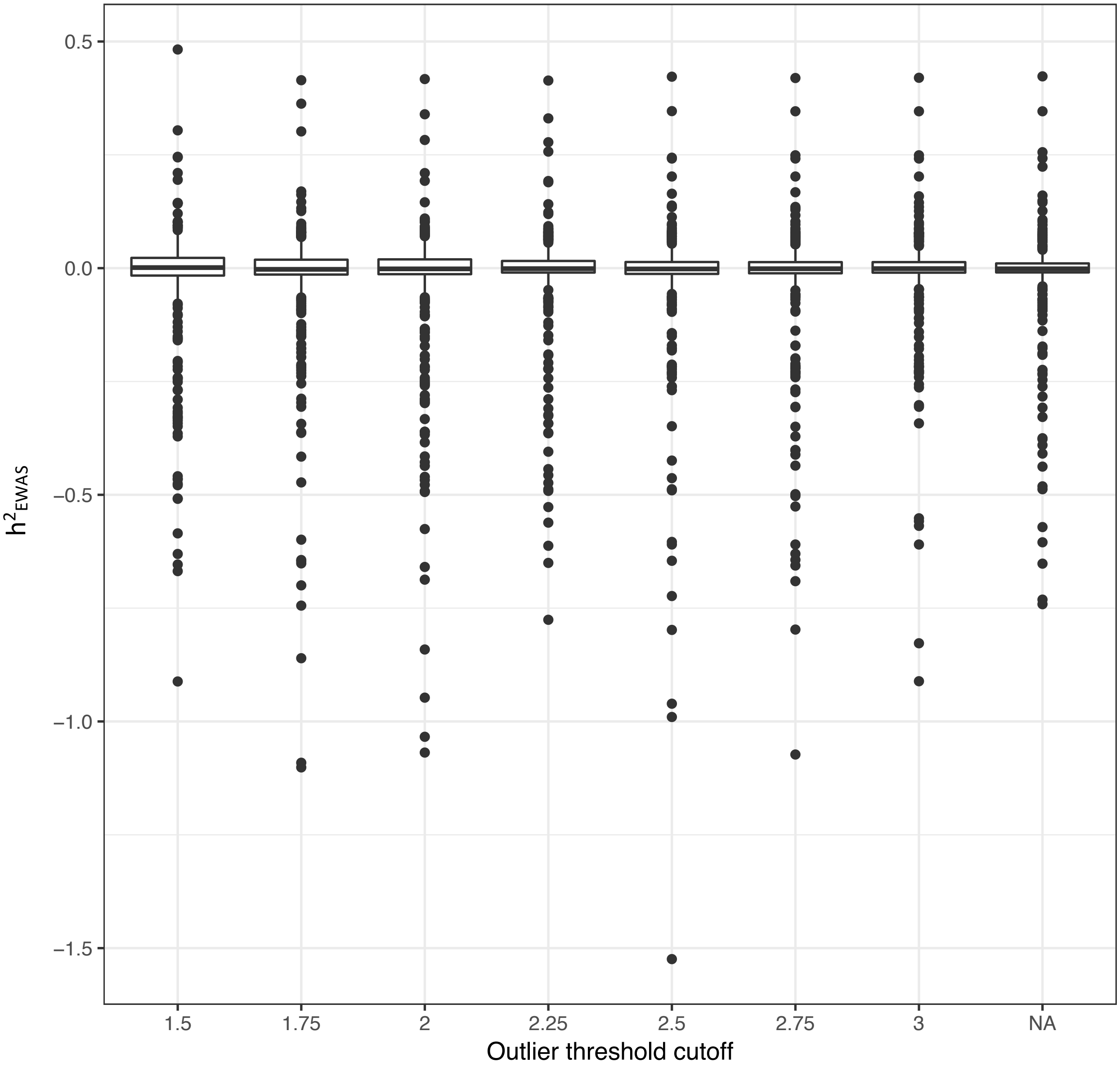


**Supplementary figure 4** Association between differing methylation kinship matrix diagonal outlier cutoff thresholds and $h^2_{EWAS}$. NA = no individuals excluded from the analysis.

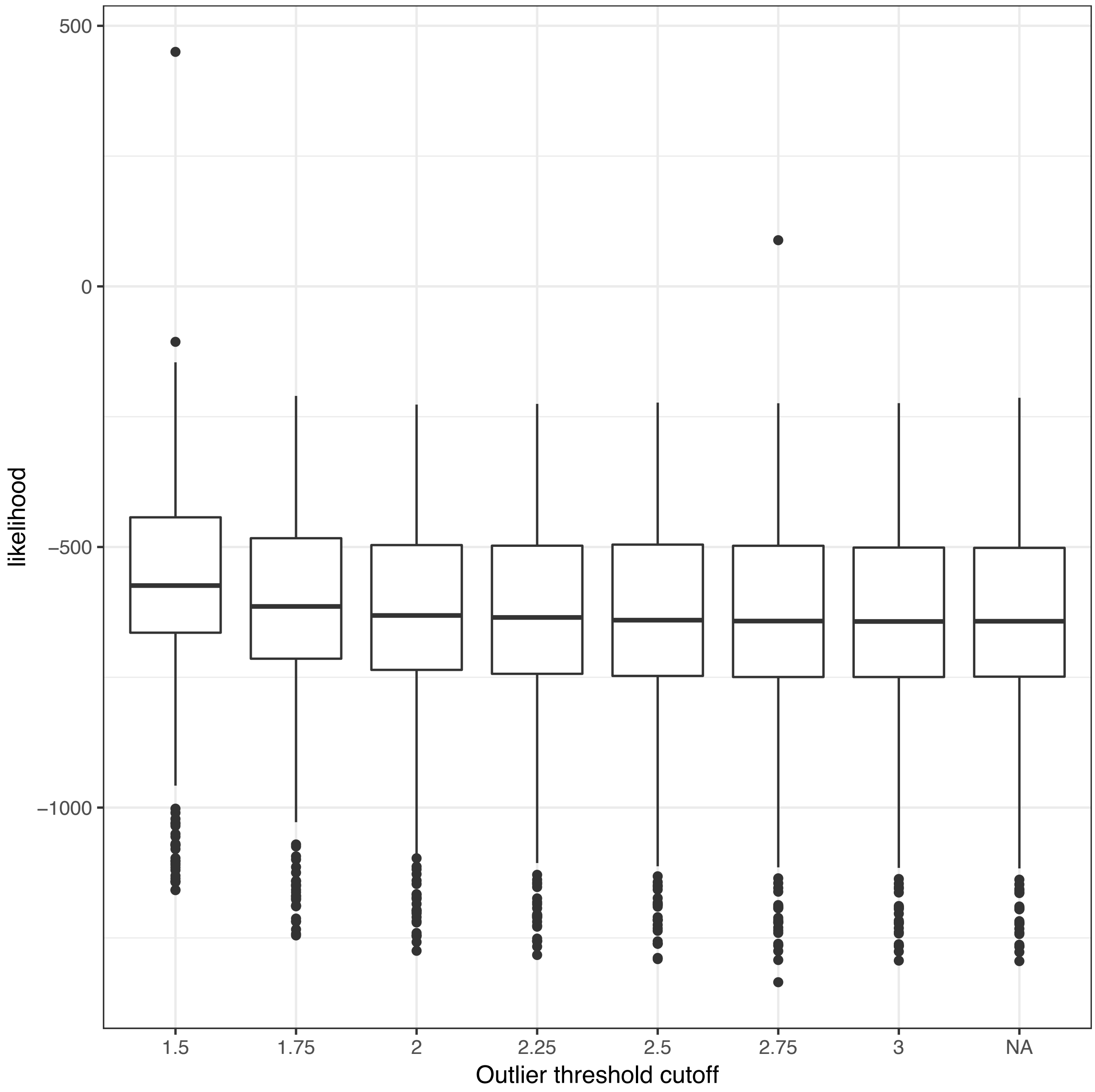


**Supplementary figure 5** How the likelihood of the REML models varied when removing individuals that had diagonal values of within the methylation relationship matrix above a certain threshold from the matrix. NA = no individuals excluded from the analysis.

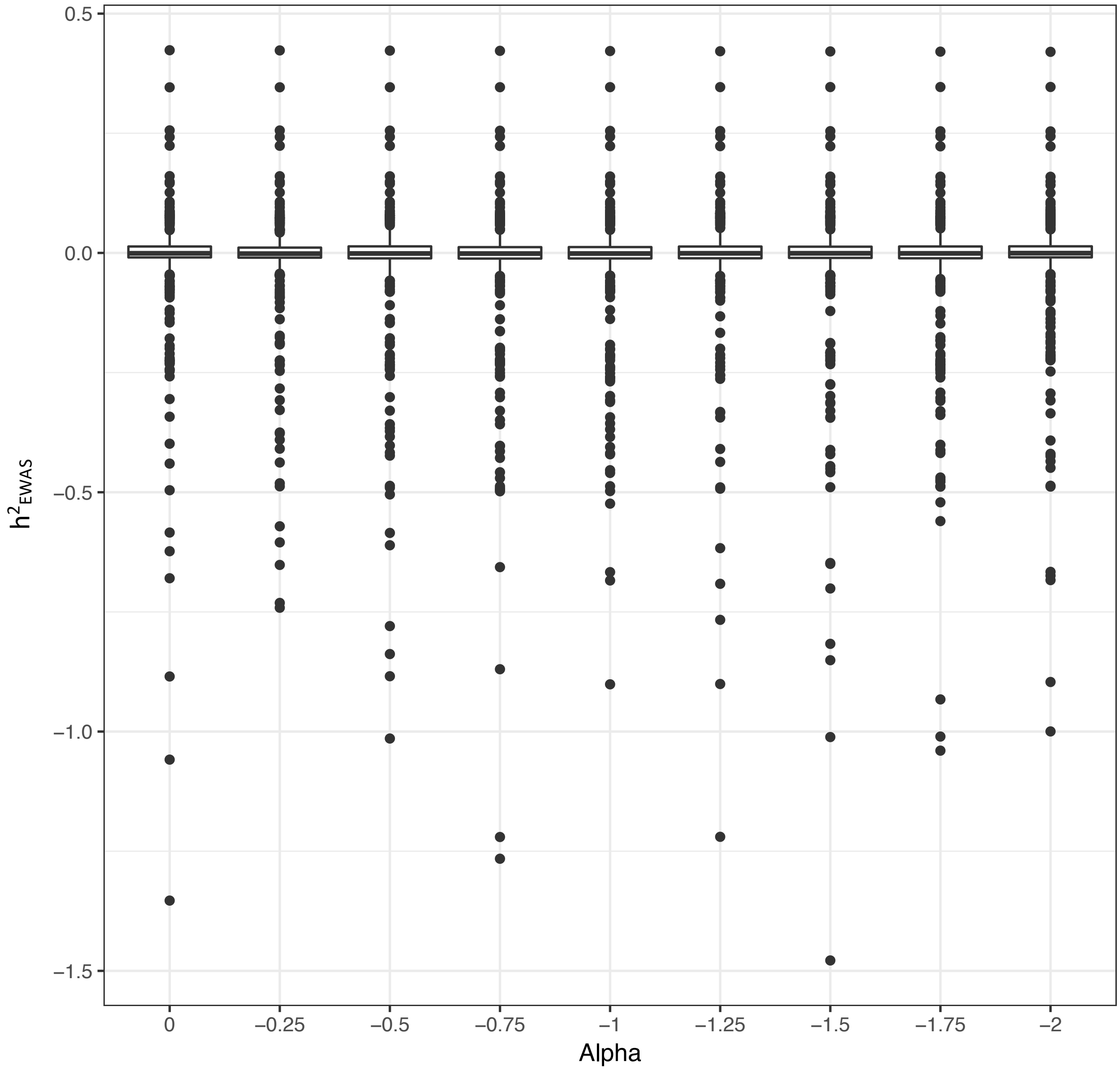


**Supplementary figure 6** Association between differing alpha values (weighting of predictors when producing the MRM) and $h^2_{EWAS}$

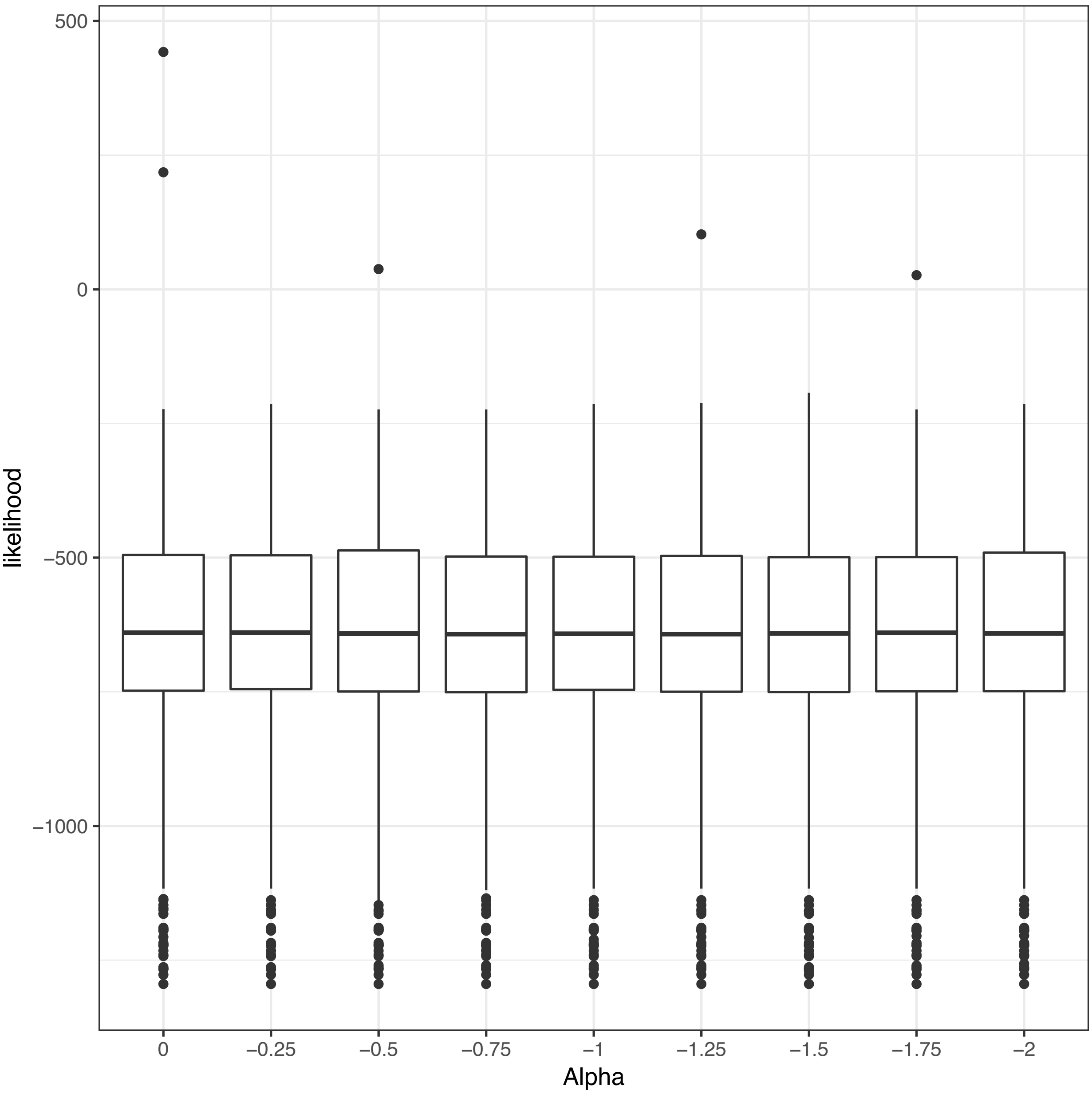


**Supplementary figure 7** The likelihood of REML models when changing the weight given to DNA methylation sites of differing variance (alpha). When alpha = -1, it is assumed that sites of high and low variance contribute equally to the $m^2$ value, as alpha increases more weight is given to sites of higher variance.

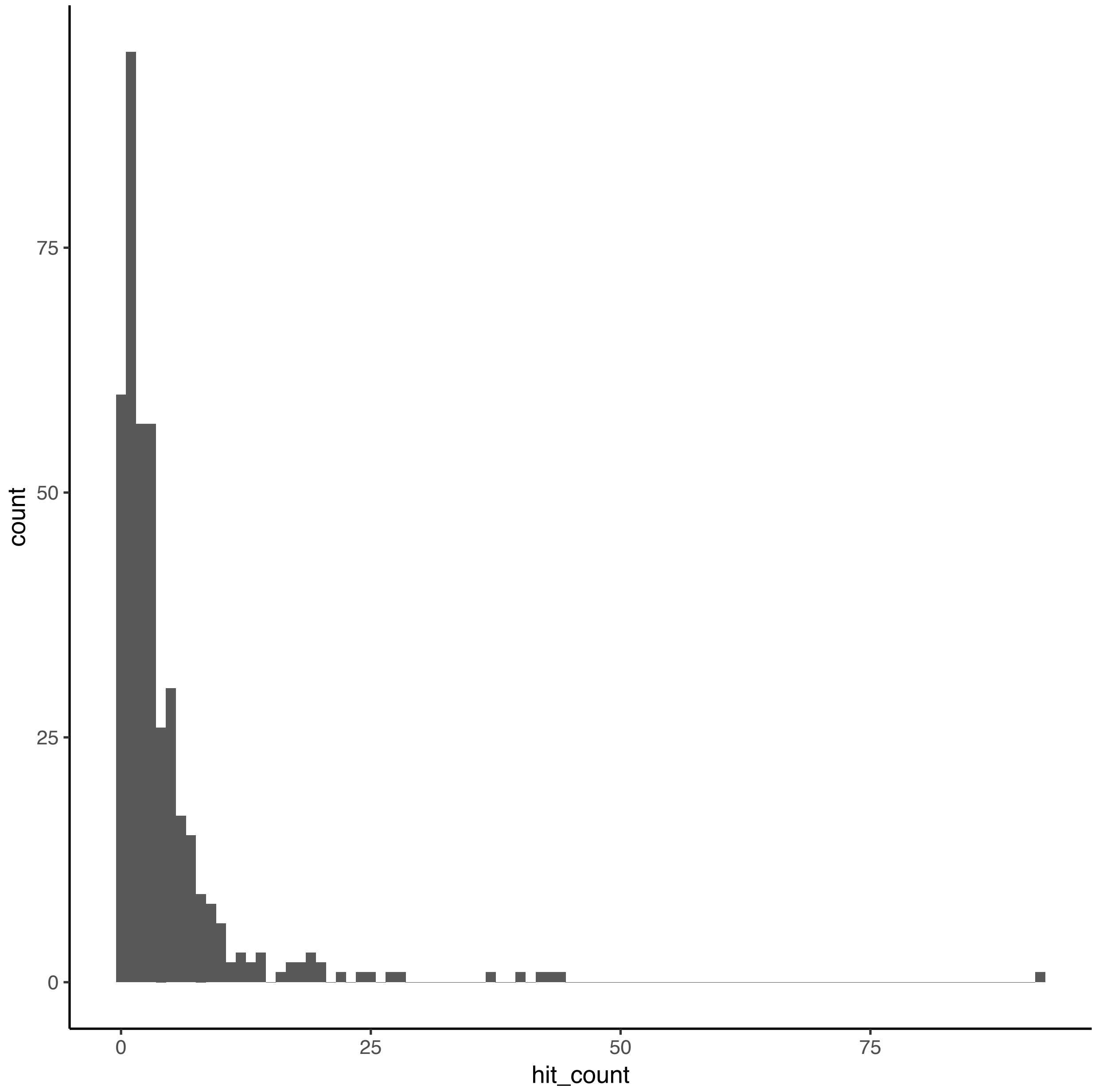


**Supplementary figure 8** The distribution of the number of DNA methylation sites identified at $P<1x10^{-5}$ across EWAS of 400 traits.

| outlier-cut-off | $h^2_{EWAS}$ summary statistics | | | | | $h^2_{EWAS} > 0$ |
|---|---|---|---|---|---|---|
| | min | q1 | med | q3 | max | |
| No cut-off | -0.741 | -0.009 | -0.001 | 0.010 | 0.423 | 13 |
| 1.5 | -0.654 | -0.013 | 0.001 | 0.020 | 0.482 | 15 |
| 1.75 | -0.860 | -0.011 | -0.002 | 0.016 | 0.415 | 15 |
| 2.25 | -0.776 | -0.009 | -0.001 | 0.014 | 0.414 | 13 |
| 2.5 | -0.961 | -0.010 | -0.001 | 0.013 | 0.422 | 15 |
| 2.75 | -1.073 | -0.009 | -0.001 | 0.013 | 0.419 | 15 |
| 2 | -0.841 | -0.009 | -0.001 | 0.017 | 0.417 | 13 |
| 3 | -0.568 | -0.009 | -0.001 | 0.013 | 0.420 | 16 |

**Supplementary table 2** Summary of $h^2_{EWAS}$ estimates for the "outlier removal" sensitivity analyses. Methylation relationship matrix diagonal values were examined and those samples that had a diagonal value above the threshold stated under the "outlier-cut-off" column were removed, and the analysis was repeated using the blanket model. $h^2_{EWAS} > 0$ = number of traits for which there was any evidence at $P<0.05$ that $h^2_{EWAS}$ was greater than 0.

| alpha | $h^2_{EWAS}$ summary statistics | | | | | $h^2_{EWAS} > 0$ |
|---|---|---|---|---|---|---|
| | min | q1 | med | q3 | max | |
| -0.25 | -0.741 | -0.010 | -0.001 | 0.011 | 0.423 | 13 |
| -0.5 | -0.838 | -0.009 | -0.001 | 0.012 | 0.423 | 13 |
| -0.75 | -0.498 | -0.009 | 0.000 | 0.013 | 0.422 | 13 |
| 0 | -0.885 | -0.008 | 0.000 | 0.012 | 0.424 | 13 |
| -1.25 | -0.617 | -0.010 | -0.001 | 0.012 | 0.421 | 13 |
| -1.5 | -0.817 | -0.009 | 0.000 | 0.013 | 0.421 | 13 |
| -1.75 | -0.560 | -0.008 | 0.000 | 0.013 | 0.421 | 13 |
| -1 | -0.684 | -0.009 | -0.001 | 0.012 | 0.422 | 13 |
| -2 | -0.896 | -0.009 | 0.000 | 0.013 | 0.420 | 13 |

**Supplementary table 3** Summary of $h^2_{EWAS}$ estimates for the "alpha" sensitivity analyses. When producing the methylation relationship matrix, probes were scaled by their observed variance and the weighting of each probe was based on the variance of DNA methylation at that site (see Methods in main manuscript for formula). Higher alpha values give more weight to sites with greater variance. $h^2_{EWAS} > 0$ = number of traits for which there was any evidence at $P<0.05$ that $h^2_{EWAS}$ was greater than 0.

| Model | Log(likelihood) | DF |
|---|---|---|
| Poisson | -1561 | 2 |
| Negative binomial | -972 | 3 |
| Hurdle-negative binomial | -954 | 5 |
| Hurdle | -1500 | 4 |
| Zero-inflated negative binomial | -972 | 5 |
| Zero-inflated Poisson | -1501 | 4 |

**Supplementary table 5** Summary of how well models fit to test the association between $h^2_{EWAS}$ and the number of differentially methylated positions identified across 400 traits at $P < 1x10^{-5}$. DF = degrees of freedom.
